# Beyond Regional Activations: Structural Connectivity Message-Passing Shallow Neural Networks for Brain Decoding

**DOI:** 10.64898/2026.03.22.713504

**Authors:** Maria Beatriz Ramos, José Diogo Marques dos Santos, Bruno Direito, Luís Paulo Reis, José Paulo Marques dos Santos

**Affiliations:** Faculty of Medicine, University of Porto, Porto, Portugal; Faculty of Engineering, University of Porto, Porto, Portugal; LIACC – Artificial Intelligence and Computer Science Laboratory, University of Porto, Portugal; CISUC – Centre for Informatics and Systems, University of Coimbra, Coimbra, Portugal; LASI – Intelligent Systems Associate Laboratory, Guimarães, Portugal; University of Maia, Maia, Portugal

## Abstract

Brain decoding from fMRI data using artificial neural networks traditionally operates at the regional level, identifying which brain areas activate during tasks but ignoring how these regions interact through structural networks. While Graph Neural Networks can capture connectivity, they require prohibitively large datasets for typical neuroscience studies. We introduce a message-passing mechanism that allows a shallow neural network to incorporate structural connectivity, enabling network-level interpretation from limited data. Using motor task data from 30 Human Connectome Project subjects, we evaluate seven structural connectivity matrices derived from deterministic and probabilistic tractography. Our approach achieves 83.0% classification accuracy while revealing functional network organization. We demonstrate that sparser, anatomy-driven connectivity matrices outperform dense alternatives, and that normalizing for network size improves model performance. Critically, our method is capable of exposing structural pathways contributing towards classification, distinguishing between complete network recruitment and selective regional activation. This approach bridges the gap between high-performance brain decoding and biological fidelity of the model, enhancing neuroscientific understanding, with implications for analyzing network dysfunctions in neurological disorders such as Alzheimer’s disease (AD), attention deficit hyperactivity disorder (ADHD), autism spectrum disorder (ASD), bipolar disorder, mild cognitive impairment (MCI), and schizophrenia.

## 1 Introduction

Artificial neural networks (ANNs) are capable of extracting valuable information from data, producing models with high predictive accuracy when properly trained and strong generalization performance when the input data are sufficiently representative [Haykin, 2009]. Functional magnetic resonance imaging (fMRI) is a neuroscientific technique that generates four-dimensional maps of brain function and has applications ranging from mental health research to brain decoding, aiming to improve our understanding of the human mind. Although early work has explored the use of ANNs for the analysis of fMRI data [Marques dos Santos, et al., 2014; Santos and Moutinho, 2011], methodological challenges have limited their broader applicability. Reviews of machine learning approaches for fMRI classification have highlighted specific concerns regarding the use of ANNs in this context [Pereira, et al., 2009]. More recently, these concerns and methodological issues are addressed through approaches explicitly designed to account for the specific characteristics of fMRI data, leading to models and results that are validated against established neuroscientific knowledge [Marques dos Santos and Marques dos Santos, 2023; Marques dos Santos and Marques dos Santos, 2022]. The developed procedures have successfully been applied to cognitive paradigms [Marques dos Santos, et al., 2025; Marques dos Santos and Marques dos Santos, 2024].

However, it focuses on a region-level analysis, only revealing which regions are recruited for the stimulus response. In mental disorders such as Alzheimer’s disease (AD), attention deficit hyperactivity disorder (ADHD), autism spectrum disorder (ASD), bipolar disorder, mild cognitive impairment (MCI), and schizophrenia, it is critical to obtain not only this information but also how these regions are organized into networks interacting with each other [Du, et al., 2018]. To achieve this aim, it is necessary to include network-level structural connectivity information either in the model architecture or in the inputs. In other domains, a common method is to leverage Graph Neural Networks (GNNs), since these can tackle the non-Euclidean properties of these complex networks. GNNs have been applied to fMRI analysis, but such approaches have focused on using functional connectivity [Li, et al., 2021]. Unlike structural connectivity, which reflects biological reality in the human brain, functional connectivity can lead to spurious conclusions, as such restrictions are not imposed. All that is measured is the statistical similarity of the signal across pairs of regions. Furthermore, due to their deep architecture, GNNs may be too complex to train on data from a typical study, around 30 subjects.

Although no major neuroscientific advances in motor control are anticipated, simple experimental paradigms remain critical for developing and validating new analytical methods. In this context, the motor paradigm adopted in this study offers a controlled, well-characterized task with well-defined regions of interest, while preserving the intrinsic challenges of fMRI data analysis. This balance facilitates the identification of methodological strengths and limitations and increases the likelihood that the proposed approach generalizes to more complex cognitive paradigms.

Thus, in this study, we aim to leverage a message-passing mechanism similar to the one present in GNNs but applied to a simpler Shallow Neural Network (SNN). The research questions are then:

- Can Shallow Neural Networks translate their good performance from region-level to network-level analysis?
- What is the method of obtaining the structural connectivity that works best when paired with the message passing mechanism and the SNN classifier?

## 2 Methods

### 2.1 Dataset, Subjects, and Experimental Paradigm

The dataset comprises 30 subjects from the Human Connetome Project (HCP) Young Adult database (https://www.humanconnectome.org/study/hcp-young-adult), specifically subjects 100307 through 124422, drawn from the 100 Unrelated Subjects subset [Elam, et al., 2021; Van Essen and Glasser, 2016; Van Essen, et al., 2013]. The subjects are randomly divided into training and testing cohorts, consisting of 20 and 10 subjects, respectively. The motor task paradigm includes five classes: left foot movement (LF), left-hand digit flexion (LH), right foot movement (RF), right-hand digit flexion (RH), and tongue movement (T). Because different acquisition sequences are used across the two sessions, each session is analyzed independently. Consequently, the training cohort comprises 40 data files (20 subjects × 2 sessions), while the testing cohort comprises 20 data files (10 subjects × 2 sessions), with no overlap between the two cohorts.

### 2.2 Feature Extraction

To extract features for training the SNN (Shallow Neural Network), the brain is parcellated according to the HCP-MMP1.0 (Human Connectome Project—multimodal parcellation) [Glasser, et al., 2016] atlas. Mean regional time series are obtained by masking each subject’s blood oxygen level–dependent (BOLD) data with every atlas region and averaging the voxel-wise signals within each region. This procedure yields 360 regional time series per session, each representing the BOLD signal from a specific brain region throughout the entire session.

Following time-series extraction, the feature selection and data standardization are performed. The regional time series are first segmented according to the stimulus sequence, and the seventh, eighth, and ninth time points following stimulus onset are averaged to form a single feature. This feature represents the class-specific neural response for each brain region. The training and testing input matrices are then constructed independently and standardized across each entire matrix. Both matrices have 360 columns corresponding to brain regions but differ in the number of samples: the training matrix has 400 instances, whereas the testing matrix has 200.

### 2.3 Structural Connectivity Matrices

The structural connectivity matrices are constructed from previously published, publicly available structural connectomes that use the HCP-MMP1.0 cortical parcellation. Three connectomes are selected, each derived using a distinct tractography strategy: (i) the *anatomy-driven deterministic connectivity matrix* proposed by Baker et al. [2018], (ii) the *population-based deterministic connectivity matrix* developed by Yeh et al. [2018], and (iii) the *probabilistic connectivity matrix* described by Rosen et al. [2018].

The anatomy-driven deterministic connectivity matrix is derived from deterministic tractography of diffusion-weighted images from 10 subjects using Generalized Q-Sampling Imaging (ratio = 1.25; angular threshold = 45°) with manual seeding targeting well-established anatomical pathways. Connectivity is defined by the presence of streamlines between regions of the HCP-MMP parcellation, yielding a group-averaged binary matrix comprising 2,100 connections. A more detailed explanation of this process is available in the original studies [Briggs, et al., 2018; Briggs, et al., 2018; Briggs, et al., 2018; Briggs, et al., 2018; Briggs, et al., 2018; Conner, et al., 2018; Conner, et al., 2018; Conner, et al., 2018; Sali, et al., 2018].

The population-based deterministic connectivity matrix is constructed from 1,065 subjects using deterministic tractography with randomized tracking parameters (GQI ratio = 1.7; angular thresholds = 15°–90°) and automated seeding based on templates of 52 major white matter bundles. Connectivity is originally defined as the relationship between white matter tracts and cortical regions and is expressed as a population-based connection probability. A more detailed explanation of this process is available in the original study [Yeh, 2022]. To obtain binary region-by-region connectivity matrices, probability thresholds of 5% (most broad), 70% (intermediate), and 99% (most restrictive) are applied, and pairs of cortical regions are considered structurally connected if both regions exhibited connection probabilities exceeding the selected threshold with the same white matter tract.

The probabilistic connectivity matrix is derived from probabilistic tractography (FSL probtrackx^1^) applied to diffusion data from 1,065 subjects, with voxel-to-parcel seeding. Inter-regional connectivity is quantified by the log-transformed number of streamlines. A more detailed explanation of this process is available in the original study [Rosen and Halgren, 2021]. As previously done to obtain binary connectivity matrices, the group-averaged matrices are thresholded at −4 (most broad), −3 (intermediate), and −2 (most restrictive). Since connectivity is measured in a logarithmic scale with base 10, the threshold −4 corresponds to 10^−4^.

For all cases, a diagonal matrix is added to the connectivity matrix. Since the connectivity matrices show only the subset of regions a given region is connected to, this ensures that, when the message-passing mechanism is applied, the signal from the region around the network is centered is still considered, i.e., the diagonal represents intra-regional connectivity. Figure 1 shows the cord graphs for all the connectivity matrices used. This representation helps visualize the difference in sparsity between the broadest and the most restrictive matrices.

**Figure 1.**
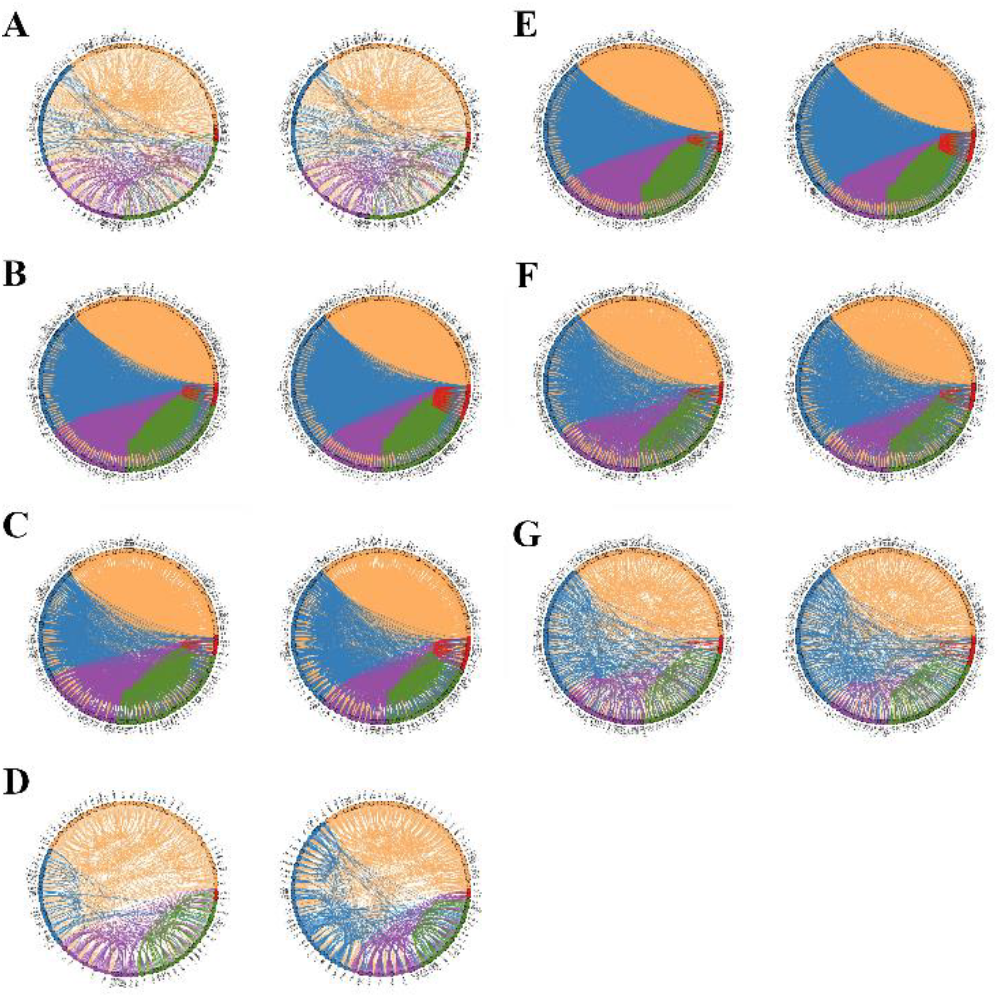
Cord graphs for all connectivity matrices. Orange connections originate in the Frontal cortex, blue in the Parietal, Purple in the Occipital, green in the Temporal, red in the Insula, and brown in the Limbic system. Pane A is the anatomy-driven deterministic connectivity matrix; panes B, C, and D are the population-based deterministic connectivity matrices, thresholds 5 %, 70%, and 99 %, respectively; panes E, F, and G are the probabilistic connectivity matrices, thresholds -4, -3, and -2, respectively. Left hemisphere is on the left side and right hemisphere on the right side for each pane.

### 2.4 Overlap Between Connectivity Matrices

To quantify the degree of overlap among the seven connectivity matrices, the Jaccard index is computed for all possible matrix pairs, according to the following definition:

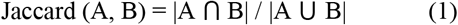

Higher Jaccard index values indicate a greater proportion of shared connections between matrices A and B. To complement this analysis, the overlap coefficient is also calculated for each pair of matrices, defined as:

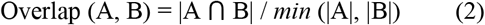

The overlap coefficient quantifies the extent to which the connections of the smaller matrix are contained within the larger one. An overlap value of 1 indicates that all connections present in the smaller matrix are also present in the other matrix.

### 2.5 Message-Passing Mechanism

The message-passing mechanism constitutes the functional foundation of graph neural networks (GNNs). It iteratively updates each node’s value based on its neighbors’ values. In the present work, only a single message-passing step is applied. Mathematically, the implementation consists of multiplying the signal matrices (training and testing) by each structural connectivity matrix. Because the diagonal is added to the structural connectivity matrices, the updated signal of each brain region corresponds to the sum of its own signal and the signals of all regions connected to it. This follows the equation:

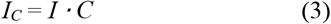

where *I* is the input matrix yielded by the feature extraction step (n×360 matrix, where n = 400 for training and n = 200 for testing), *C* is the structural connectivity matrix (360×360 matrix) and *I*_*C*_ is the input matrix after one step of the message passing mechanism according to the structural connectivity matrix *C* (n×360 matrix, where n = 400 for training and n = 200 for testing).

A correction factor is also evaluated in all cases. This factor consists of dividing the summed signal by the total number of connected regions in the network (i.e., the sum over all non-zero values in the relevant column of matrix *C*), thereby yielding an average network signal. This correction is introduced to mitigate the expected problem of signal dilution, which occurs when not all regions of the network are recruited in response to a given stimulus. In such cases, summing activation-related signal peaks with baseline activity reduces the distinction between activated and resting networks. The proposed correction factor is therefore expected to improve the performance of connectivity matrices that represent more extensive or encompassing networks.

### 2.6 SNN Training, Pruning, and Retraining

To keep consistency with other studies, the fully connected SNN consists of 360 input nodes (one per brain region or network), 10 hidden nodes in a single hidden layer, and 5 output nodes (one per class in the paradigm design).

The network architecture is implemented through the AMORE library, version 0.2-15 [Limas, et al., 2014]; R, version 4.5.1; and RStudio, version 2025.05.1.513. The activation function used for the hidden nodes is the “tansig” (hyperbolic tangent). The tansig outputs values in the ]−1, 1[ range, with the function centered at 0. The activation function for the output nodes is “sigmoid”, which outputs values in the ]0, 1[ range.

During training, a grid search is performed to find the best hyperparameters (learning rate and momentum). During the grid search, each learning rate and momentum pair is used 100 times to account for variations in the randomness of weight initialization. Afterward, the best learning rate and momentum pair are used to train 50,000 networks, aiming to achieve the best possible network. In this step, the only variation between networks is the randomly initialized weights. This step yields the “fully connected network”.

The first stage of the pruning process consists of ranking all possible path-weights. The concept of “path-weights” [Marques dos Santos and Marques dos Santos, 2023; Marques dos Santos and Marques dos Santos, 2022], considers the weights of all connections along the path from one input to one output, passing through one hidden node (because the network architecture in this study involves only one hidden layer). Thus, in mathematical notation, a path-weight is as follows:

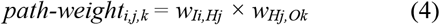

where *w*_*Ii,Hj*_ is the weight between the input node *Ii* and the hidden node *Hj*; and *w*_*Hj,Ok*_ is the weight between the hidden node *Hj* and the output node *Ok*. Therefore, *path-weight*_*i,j*,_*k* is the product of the weights along the path from input *Ii* to output *Ok*, passing by the hidden node *Hj*. A path-weight near zero tends to nullify the input signal in the path to the output, and, therefore, it may be pruned, as it is of low relevance. On the contrary, a path-weight away from zero, whether positive or negative, is more relevant for classification and, therefore, must be retained in the network.

All series of path-weights have two elbows, meaning that few paths leverage the information from the input to influence the model’s output. The elbow points are computed separately for the positive and negative parts, using the “pathviewer” package for R [Baliga, et al., 2023]. After ranking and plotting all the inputs, the path-weights falling between the positive and negative elbows are pruned. This process is explained in further detail elsewhere [Marques dos Santos and Marques dos Santos, 2024].

The “pruned network” is then retrained to update the weights to the new architecture. During the process, a grid search for the best hyperparameters is also performed. However, only one network is trained per pair, since there is no randomness in the initial weights. The intent behind the retraining process is to allow the network to recalculate its weights within the new (lighter) architecture, expecting it to regain the accuracy lost during the pruning process. This new training step is expected to be more efficient than the initial one, as the irrelevant network connections have been removed. This step yields the “retrained network”.

## 3 Results

### 3.1 Overlap Between Connectivity Matrices

Overall, the Jaccard index values (Table 1) are consistently low across all pairwise comparisons, indicating limited overlap between the connectivity matrices. This pattern is observed both across matrices derived from different tractography approaches and among matrices obtained from different threshold levels within the same connectome. In the latter case, low Jaccard values arise despite the nested nature of the thresholded matrices, as increasing the threshold primarily enlarges the union of connections while the intersection remains unchanged.

**Table 1.** Jaccard Indexes, “Anatomical” indicates the anatomy-driven deterministic connectivity matrix, “Pop.” indicates the population-based deterministic connectivity matrices followed by the respective threshold, and “Prob.” indicates the probabilistic connectivity matrices followed by the respective threshold.

| Jaccard | Anatomical | Pop. 5% | Pop. 70% | Pop. 99% | Prob. (-4) | Prob. (-3) | Prob. (-2) |
| --- | --- | --- | --- | --- | --- | --- | --- |
| Number of Connections | 2,100 | 36,942 | 14,258 | 2,732 | 65,790 | 16,750 | 3,762 |
| Anatomical |  | 0.0525 | 0.101 | 0.193 | 0.024 | 0.039 | 0.077 |
| Pop. 5% |  |  | 0.385 | 0.073 | 0.342 | 0.179 | 0.065 |
| Pop. 70% |  |  |  | 0.191 | 0.141 | 0.124 | 0.079 |
| Pop. 99% |  |  |  |  | 0.031 | 0.054 | 0.099 |
| Prob. (-4) |  |  |  |  |  | 0.254 | 0.057 |
| Prob. (-3) |  |  |  |  |  |  | 0.224 |
| Prob. (-2) |  |  |  |  |  |  |  |

Differences in matrix size further modulate Jaccard index values, with smaller indices observed for comparisons involving matrices with markedly different numbers of connections. This effect is particularly evident when contrasting sparse matrices with substantially denser ones, where the union of connections increases considerably while the shared subset remains relatively small. Nevertheless, comparisons between matrices derived using deterministic tractography methods tend to yield higher Jaccard indices than comparisons involving deterministic and probabilistic matrices, suggesting greater similarity among deterministic approaches.

The overlap coefficient (Table 2) reveals a complementary pattern. Comparisons between different threshold levels within the population-based deterministic connectivity and probabilistic connectivity yield overlap values of one, reflecting the fact that these matrices represent progressively filtered versions of the same original connectome and that all connections present in the sparser matrix are also contained in the denser one. In contrast, comparisons across different connectomes show overlap coefficients that vary with relative matrix density: as the number of connections in one matrix decreases, the overlap coefficient with a denser matrix also decreases, indicating that fewer connections are shared.

**Table 2.** Overlap Coefficients, “Anatomical” indicates the anatomy-driven deterministic connectivity matrix, “Pop.” indicates the population-based deterministic connectivity matrices followed by the respective threshold, and “Prob.” indicates the probabilistic connectivity matrices followed by the respective threshold.

| Overlap | Anatomical | Pop. 5% | Pop. 70% | Pop. 99% | Prob. (-4) | Prob. (-3) | Prob. (-2) |
| --- | --- | --- | --- | --- | --- | --- | --- |
| Number of Connections | 2,100 | 36,942 | 14,258 | 2,732 | 65,790 | 16,750 | 3,762 |
| Anatomical |  | 0.928 | 0.719 | 0.373 | 0.758 | 0.343 | 0.200 |
| Pop. 5% |  |  | 1.000 | 1.000 | 0.708 | 0.488 | 0.669 |
| Pop. 70% |  |  |  | 1.000 | 0.697 | 0.240 | 0.354 |
| Pop. 99% |  |  |  |  | 0.757 | 0.366 | 0.215 |
| Prob. (-4) |  |  |  |  |  | 1.000 | 1.000 |
| Prob. (-3) |  |  |  |  |  |  | 1.000 |
| Prob. (-2) |  |  |  |  |  |  |  |

When matrices have similar connection densities, overlap coefficients are consistently lower, further indicating that the three connectomes represent distinct sets of structural connections. Among the cross-connectome comparisons, the greatest degree of overlap is observed between the matrices derived from population-based deterministic connectivity and the anatomy-driven deterministic connectivity, with the densest population-based deterministic connectivity matrix encompassing approximately 93% of the connections present in the anatomy-driven deterministic connectivity matrix.

### 3.2 Region-Level Classification

For the region-level classification, each of the 360 inputs corresponds to the signal of only one region in the HCP-MMP1.0 atlas. The accuracies for all network stages are present in Table 3. The fully connected network achieves 90.0 % accuracy, dropping to 83.0 % with pruning, but partially recovers to 85.0 % with retraining. Looking at partial accuracies, the model performs worse for the LF class across all three stages. As shown in the confusion matrix (cf. Supplementary Materials), the most common error for LF instances is the classifier assigning the RF output, indicating that the model has the most trouble separating between the two feet. Pruning negatively affects the identification of RF and RH instances primarily, and, interestingly, has a slight positive effect on the LF class. The mitigation of these effects during retraining is what leads to the recovery of lost accuracy.

**Table 3.**
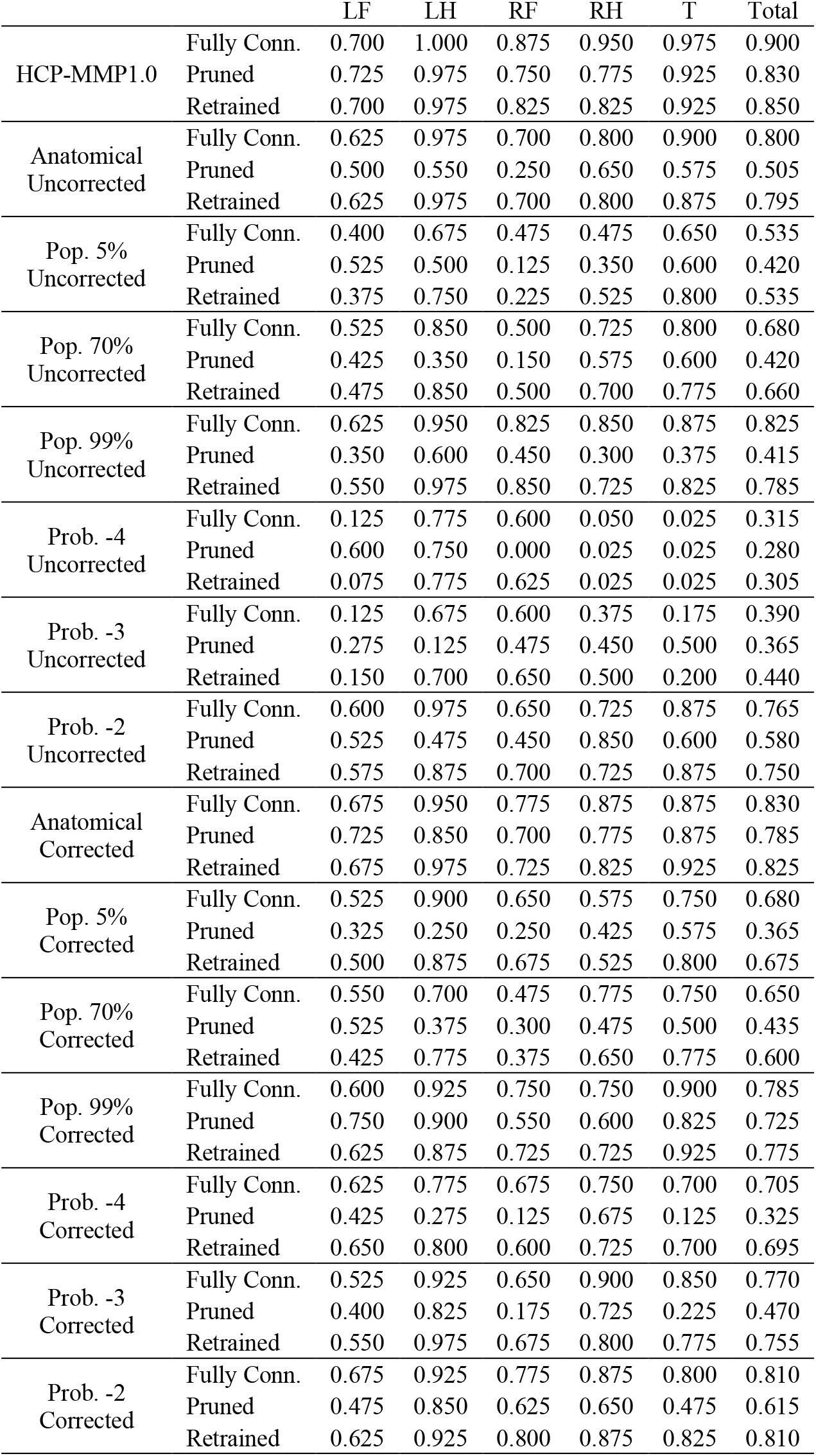
Global and partial accuracies of the full, pruned, and retrained networks for all classifiers, “Anatomical” indicates the anatomy-driven deterministic connectivity matrix, “Pop.” indicates the population-based deterministic connectivity matrices followed by the respective threshold, and “Prob.” indicates the probabilistic connectivity matrices followed by the respective threshold.

|  |  | LF | LH | RF | RH | T | Total |
| --- | --- | --- | --- | --- | --- | --- | --- |
| HCP-MMP1.0 | Fully Conn. | 0.700 | 1.000 | 0.875 | 0.950 | 0.975 | 0.900 |
|  | Pruned | 0.725 | 0.975 | 0.750 | 0.775 | 0.925 | 0.830 |
|  | Retrained | 0.700 | 0.975 | 0.825 | 0.825 | 0.925 | 0.850 |
| Anatomical<br>Uncorrected | Fully Conn. | 0.625 | 0.975 | 0.700 | 0.800 | 0.900 | 0.800 |
|  | Pruned | 0.500 | 0.550 | 0.250 | 0.650 | 0.575 | 0.505 |
|  | Retrained | 0.625 | 0.975 | 0.700 | 0.800 | 0.875 | 0.795 |
| Pop. 5%<br>Uncorrected | Fully Conn. | 0.400 | 0.675 | 0.475 | 0.475 | 0.650 | 0.535 |
|  | Pruned | 0.525 | 0.500 | 0.125 | 0.350 | 0.600 | 0.420 |
|  | Retrained | 0.375 | 0.750 | 0.225 | 0.525 | 0.800 | 0.535 |
| Pop. 70%<br>Uncorrected | Fully Conn. | 0.525 | 0.850 | 0.500 | 0.725 | 0.800 | 0.680 |
|  | Pruned | 0.425 | 0.350 | 0.150 | 0.575 | 0.600 | 0.420 |
|  | Retrained | 0.475 | 0.850 | 0.500 | 0.700 | 0.775 | 0.660 |
| Pop. 99%<br>Uncorrected | Fully Conn. | 0.625 | 0.950 | 0.825 | 0.850 | 0.875 | 0.825 |
|  | Pruned | 0.350 | 0.600 | 0.450 | 0.300 | 0.375 | 0.415 |
|  | Retrained | 0.550 | 0.975 | 0.850 | 0.725 | 0.825 | 0.785 |
| Prob. -4<br>Uncorrected | Fully Conn. | 0.125 | 0.775 | 0.600 | 0.050 | 0.025 | 0.315 |
|  | Pruned | 0.600 | 0.750 | 0.000 | 0.025 | 0.025 | 0.280 |
|  | Retrained | 0.075 | 0.775 | 0.625 | 0.025 | 0.025 | 0.305 |
| Prob. -3<br>Uncorrected | Fully Conn. | 0.125 | 0.675 | 0.600 | 0.375 | 0.175 | 0.390 |
|  | Pruned | 0.275 | 0.125 | 0.475 | 0.450 | 0.500 | 0.365 |
|  | Retrained | 0.150 | 0.700 | 0.650 | 0.500 | 0.200 | 0.440 |
| Prob. -2<br>Uncorrected | Fully Conn. | 0.600 | 0.975 | 0.650 | 0.725 | 0.875 | 0.765 |
|  | Pruned | 0.525 | 0.475 | 0.450 | 0.850 | 0.600 | 0.580 |
|  | Retrained | 0.575 | 0.875 | 0.700 | 0.725 | 0.875 | 0.750 |
| Anatomical<br>Corrected | Fully Conn. | 0.675 | 0.950 | 0.775 | 0.875 | 0.875 | 0.830 |
|  | Pruned | 0.725 | 0.850 | 0.700 | 0.775 | 0.875 | 0.785 |
|  | Retrained | 0.675 | 0.975 | 0.725 | 0.825 | 0.925 | 0.825 |
| Pop. 5%<br>Corrected | Fully Conn. | 0.525 | 0.900 | 0.650 | 0.575 | 0.750 | 0.680 |
|  | Pruned | 0.325 | 0.250 | 0.250 | 0.425 | 0.575 | 0.365 |
|  | Retrained | 0.500 | 0.875 | 0.675 | 0.525 | 0.800 | 0.675 |
| Pop. 70%<br>Corrected | Fully Conn. | 0.550 | 0.700 | 0.475 | 0.775 | 0.750 | 0.650 |
|  | Pruned | 0.525 | 0.375 | 0.300 | 0.475 | 0.500 | 0.435 |
|  | Retrained | 0.425 | 0.775 | 0.375 | 0.650 | 0.775 | 0.600 |
| Pop. 99%<br>Corrected | Fully Conn. | 0.600 | 0.925 | 0.750 | 0.750 | 0.900 | 0.785 |
|  | Pruned | 0.750 | 0.900 | 0.550 | 0.600 | 0.825 | 0.725 |
|  | Retrained | 0.625 | 0.875 | 0.725 | 0.725 | 0.925 | 0.775 |
| Prob. -4<br>Corrected | Fully Conn. | 0.625 | 0.775 | 0.675 | 0.750 | 0.700 | 0.705 |
|  | Pruned | 0.425 | 0.275 | 0.125 | 0.675 | 0.125 | 0.325 |
|  | Retrained | 0.650 | 0.800 | 0.600 | 0.725 | 0.700 | 0.695 |
| Prob. -3<br>Corrected | Fully Conn. | 0.525 | 0.925 | 0.650 | 0.900 | 0.850 | 0.770 |
|  | Pruned | 0.400 | 0.825 | 0.175 | 0.725 | 0.225 | 0.470 |
|  | Retrained | 0.550 | 0.975 | 0.675 | 0.800 | 0.775 | 0.755 |
| Prob. -2<br>Corrected | Fully Conn. | 0.675 | 0.925 | 0.775 | 0.875 | 0.800 | 0.810 |
|  | Pruned | 0.475 | 0.850 | 0.625 | 0.650 | 0.475 | 0.615 |
|  | Retrained | 0.625 | 0.925 | 0.800 | 0.875 | 0.825 | 0.810 |

### 3.3 Network-Level Classification

Table 3 shows the accuracy of models trained on data with a message-passing mechanism using the seven uncorrected connectivity matrices.

For the uncorrected anatomy-driven deterministic connectivity matrix, the fully connected network achieves 80.0 % accuracy, dropping to 50.5 % with pruning, and almost fully recovers, reaching 79.5 % after retraining. Here, interestingly, pruning has a particularly strong effect on LH and RF classification, with the worst-performing class in the pruned network being RF.

For the uncorrected population-based deterministic connectivity matrices, the connectivity matrix with a 5 % threshold (most broad) achieves 53.5 % with the fully connected network, dropping to 42.0 % with pruning, and fully recovers the lost performance with the retraining step (53.5 % accuracy). It is worth noting that, in this case, the RF class barely improves with retraining, and it is the worst-performing class in the final retrained network.

The connectivity matrix with a threshold of 70 % (intermediate) achieves 68.0 % with the fully connected network, dropping to 42.0 % with pruning and recovering most of the lost performance with the retraining step to 66.0 % accuracy.

The connectivity matrix with a threshold of 99 % (most restrictive) achieves 82.5 % with the fully connected network, dropping to 41.5 % with pruning and recovering most of the lost performance with the retraining step to 78.5 % accuracy. For the uncorrected probabilistic connectivity matrix. The connectivity matrix thresholded at −4 (most broad) achieves 31.5 % with the fully connected network, dropping to 28.0 % with pruning, and recovers most of the lost performance during the retraining step to 30.5 % accuracy.

The connectivity matrix thresholded at −3 (intermediate) achieves 39.0 % with the fully connected network, dropping to 36.5 % with pruning. Interestingly, for this case, the retrained network outperforms the initial fully connected network, achieving 44.0 % accuracy.

The connectivity matrix thresholded at −2 (most restrictive) achieves 76.5 % with the fully connected network, dropping to 58.0 % with pruning, and recovers most of the lost performance during the retraining step to 75.0 % accuracy.

The next paragraphs consider the models trained on data with a message-passing mechanism using the seven corrected connectivity matrices.

For the corrected anatomy-driven deterministic connectivity matrix, the fully connected network achieves 83.0 % accuracy, dropping to 78.5 % with pruning, and almost fully recovers, reaching 82.5 % after retraining. The pruning step has a similar effect on partial accuracies as for the uncorrected version of the matrix.

For the corrected population-based deterministic connectivity matrices, the connectivity matrix with a threshold of 5 % (most broad) achieves 68.0 % with the fully connected network, dropping to 36.5 % with pruning and, with the retraining step, recovers almost all of the lost performance to 67.5 % accuracy.

The connectivity matrix with a threshold of 70 % (intermediate) achieves 65.0 % with the fully connected network, dropping to 43.5 % with pruning and recovering most of the lost performance with the retraining step to 60.0 % accuracy.

The connectivity matrix with a threshold of 99 % (most restrictive) achieves 78.5 % with the fully connected network, dropping to 72.5 % with pruning and recovering most of the lost performance with the retraining step to 77.5 % accuracy. For the corrected probabilistic connectivity matrix. The connectivity matrix thresholded at −4 (most broad) achieves 70.5 % with the fully connected network, dropping to 32.5 % with pruning, and recovers most of the lost performance during the retraining step to 69.5 % accuracy.

The connectivity matrix thresholded at −3 (intermediate) achieves 77.0 % with the fully connected network, dropping to 47.0 % with pruning, and recovers most of the lost performance during the retraining step to 75.5 % accuracy.

The connectivity matrix thresholded at −2 (most restrictive) achieves 81.0 % with the fully connected network, dropping to 61.5 % with pruning, and fully recovers the lost performance during the retraining step (81.0 % accuracy).

In sum, across global accuracies, the results reveal that the anatomy-driven deterministic connectivity matrix (Anatomical) outperforms the population-based deterministic (Pop.) and probabilistic (Prob.) connectivity matrices. This is true both for uncorrected and corrected connectivity matrices, and for fully connected and retrained networks. In addition, the more restrictive the connective matrix is (Pop. 5% → 70%→ 99%; Prob. -4 → -3 → -2), the higher the accuracy. Finally, there is a tendency for accuracies from corrected networks to be higher than those from uncorrected networks.

## 4 Discussion

The top five best performing connectivity matrices are the corrected anatomy-driven deterministic connectivity matrix (83.0 % accuracy), uncorrected population-based deterministic connectivity matrix thresholded at 99 % (82.5 % accuracy), corrected probabilistic connectivity thresholded at −2 (81.0 % accuracy), uncorrected anatomy-driven deterministic connectivity matrix (80.0 % accuracy), and corrected population-based deterministic connectivity matrix thresholded at 99 % (78.5 % accuracy). These results suggest that deterministic methods outperform probabilistic methods, particularly the anatomy-driven deterministic connectivity, which appears twice in the top five.

It is also clear that sparser connectivity matrices yield better-performing classifiers, and none of the message-passing models achieve the performance of the region-based model. This may be due to signal dilution, which occurs when regions with a peak in the BOLD signal are summed with regions in the baseline. This process may make the signal peaks less explicit and, thus, harder for the classifiers to detect. Therefore, the results indicate a trade-off between including network-level information in the model to obtain explanations of the regions’ organizations in the stimulus-response and classifier performance, which provides confidence in the validity of the extracted explanations.

When looking at fully connected networks, the correction factor provides good results for the anatomy-driven deterministic connectivity matrix, all probabilistic connectivity matrices and the broadest population-based deterministic connectivity matrix: population-based deterministic connectivity matrix thresholded at 5 % from 53.5 % to 68.0 %, probabilistic connectivity thresholded at −4 from 31.5 % to 70.5 %, thresholded at −3 from 39.0 % to 77.0 %, thresholded at −2 from 76.5 % to 81.0 %, and anatomy-driven deterministic connectivity matrix from 80.0 % to 83.0 %. However, it has a negative impact on the population-based deterministic connectivity matrix thresholded at 70 % (from 68.0 % to 65.0 %) and thresholded at 99 % (from 82.5 % to 78.5 %). When looking at the final retrained networks, this trend is kept. Thus, the correction factor seems to work particularly well for probabilistic connectivity matrices and has a slight positive impact on the anatomy-driven deterministic connectivity matrix but a negative impact on most population-based deterministic connectivity matrices. Nonetheless, overall, the correction factor may be worth applying.

Looking at the most affected partial accuracies after pruning, the uncorrected anatomy-driven deterministic connectivity matrix shows the worst performing class is RF, but in the corrected version, this margin is smaller. Interestingly, the corrected version shows a smaller drop in accuracy with pruning. For the uncorrected population-based deterministic connectivity matrices, the most affected class is RF for thresholds 5 % and 70 %, but RH for threshold 99 %. For the corrected version, LH and RF perform equally badly at the 5 % threshold. RF is still the worst-performing for 70 %, and RF is, this time, also the worst-performing for 99 %. For the population-based deterministic connectivity matrices, while for the uncorrected, the impact of the pruning increases with increasing the restrictiveness of the threshold, for the corrected, the opwposite happens. The more restrictive the matrix, the less impact pruning has.

As pruning affects connectivity matrices differently, it is possible to conclude that although some global performance metrics may be similar, the models arrive at these decisions through different processes. All models seem to suffer primarily in the feet classification, with the LF class consistently problematic across all stages of all models, suggesting that this difficulty is inherent in the data. Furthermore, the other foot-related class, RF, is also commonly among the most affected by the pruning process, indicating that the networks lack the information needed to identify this class, which is concentrated in a few paths, as is the case with other classes that retain their performance better.

This same reasoning can be applied to the progression of the global accuracy across model stages. Models that drop more during pruning have their relevant information more diluted across their architecture, while models that retain performance better have their relevant information more concentrated in important paths. Thus, the correction factor seems to push the models to concentrate more relevant information (depending on fewer inputs for classification). As the correction factor reduces the drop during pruning in the more restrictive matrices, it may be that this adjustment pushes the network to concentrate the information more in the relevant paths (depending on fewer inputs). However, as it has the opposite effect on the broader matrices, in this case, it may help these matrices achieve performances similar to the restrictive matrices, but this is achieved by using information from more sources (inputs).

The conclusions of the study are threefold: 1) the anatomy-driven deterministic connectivity matrix achieves higher accuracies than the population-based deterministic and probabilistic connectivity matrices; 2) concerning the two probabilistic connectivity matrices, the more restrictive a matrix is (i.e., with a lesser number of connections), the higher the accuracy is; and 3) the introduction of a correction factor improves the model performance. Hence, because similar accuracies are achieved, using message-passing implemented by an anatomy-driven deterministic connectivity matrix with a correction factor may illuminate the brain networks participating in motor actions, providing an added layer of understanding beyond mere identification of single brain regions recruited.

## Ethical Statement

There are no ethical issues.

## Acknowledgments

This work was financially supported by UID/00027/2025 (LIACC - Artificial Intelligence and Computer Science Laboratory; DOI https://doi.org/10.54499/UID/00027/2025), UIDB/00326/2025 and UIDP/00326/2025 (CISUC – Centre for Informatics and Systems of the University of Coimbra), and LA/P/0104/2020 (LASI - Intelligent Systems Associate Laboratory), funded by Fundação para a Ciência e a Tecnologia, I.P./ MECI through national funds. The research carried out was within the scope of José Diogo Marques dos Santos scientific grant 2025.02850.BD, and Bruno Direito CEECINST/00117/2021/CP2784/CT0002 (DOI https://doi.org/10.54499/CEEC-INST/00117/2021/CP2784/CT0002).

## Footnotes

1 https://fsl.fmrib.ox.ac.uk/fsl/docs/diffusion/probtrackx.html

## Notes

### Competing Interest Statement

The authors have declared no competing interest.

