## supplementary materials for "Beyond Regional Activations: Structural Connectivity Message-Passing Shallow Neural Networks for Brain Decoding"

Supplementary Materials: Confusion Matrices

| | | LF | LH | RF | RH | T | Total |
|---|---|---|---|---|---|---|---|
| Fully Connected Network | LF | 28 | 1 | 7 | 0 | 4 | 40 |
| | LH | 0 | 40 | 0 | 0 | 0 | 40 |
| | RF | 2 | 1 | 35 | 2 | 0 | 40 |
| | RH | 0 | 2 | 0 | 38 | 0 | 40 |
| | T | 0 | 0 | 1 | 0 | 39 | 40 |
| | Total | 30 | 44 | 43 | 40 | 43 | 200 |
| | Accuracy | 0.700 | 1.000 | 0.875 | 0.950 | 0.975 | 0.900 |
| | Precision | 0.933 | 0.909 | 0.814 | 0.950 | 0.970 | |
| Pruned Network | LF | 29 | 1 | 7 | 0 | 3 | 40 |
| | LH | 0 | 39 | 0 | 1 | 0 | 40 |
| | RF | 8 | 0 | 30 | 2 | 0 | 40 |
| | RH | 1 | 4 | 3 | 31 | 1 | 40 |
| | T | 2 | 0 | 1 | 0 | 37 | 40 |
| | Total | 40 | 44 | 41 | 34 | 41 | 200 |
| | Accuracy | 0.725 | 0.975 | 0.750 | 0.775 | 0.925 | 0.830 |
| | Precision | 0.725 | 0.886 | 0.732 | 0.912 | 0.902 | |
| Retrained Network | LF | 28 | 2 | 7 | 0 | 3 | 40 |
| | LH | 0 | 39 | 0 | 1 | 0 | 40 |
| | RF | 2 | 1 | 33 | 4 | 0 | 40 |
| | RH | 0 | 3 | 3 | 33 | 1 | 40 |
| | T | 1 | 0 | 2 | 0 | 37 | 40 |
| | Total | 31 | 45 | 45 | 38 | 41 | 200 |
| | Accuracy | 0.700 | 0.975 | 0.825 | 0.825 | 0.925 | 0.850 |
| | Precision | 0.903 | 0.867 | 0.733 | 0.868 | 0.902 | |

Table 1: Confusion matrices, accuracies, and precisions for the region-level classifier using the HCP-MMP1.0 atlas.

| | | LF | LH | RF | RH | T | Total |
|---|---|---|---|---|---|---|---|
| Fully Connected Network | LF | 25 | 2 | 9 | 2 | 2 | 40 |
| | LH | 0 | 39 | 0 | 1 | 0 | 40 |
| | RF | 8 | 1 | 28 | 3 | 0 | 40 |
| | RH | 1 | 7 | 0 | 32 | 0 | 40 |
| | T | 3 | 0 | 1 | 0 | 36 | 40 |
| | Total | 37 | 49 | 38 | 38 | 38 | 200 |
| | Accuracy | 0.625 | 0.975 | 0.700 | 0.800 | 0.900 | 0.800 |
| | Precision | 0.676 | 0.796 | 0.737 | 0.842 | 0.947 | |
| Pruned Network | LF | 20 | 5 | 6 | 6 | 3 | 40 |
| | LH | 15 | 22 | 0 | 2 | 1 | 40 |
| | RF | 10 | 3 | 10 | 16 | 1 | 40 |
| | RH | 3 | 8 | 2 | 26 | 1 | 40 |
| | T | 14 | 0 | 0 | 3 | 23 | 40 |
| | Total | 62 | 38 | 18 | 53 | 29 | 200 |
| | Accuracy | 0.500 | 0.550 | 0.250 | 0.650 | 0.575 | 0.505 |
| | Precision | 0.323 | 0.579 | 0.556 | 0.491 | 0.793 | |
| Retrained Network | LF | 25 | 2 | 9 | 3 | 1 | 40 |
| | LH | 0 | 39 | 0 | 1 | 0 | 40 |
| | RF | 6 | 1 | 28 | 5 | 0 | 40 |
| | RH | 1 | 5 | 2 | 32 | 0 | 40 |
| | T | 3 | 1 | 0 | 1 | 35 | 40 |
| | Total | 35 | 48 | 39 | 42 | 36 | 200 |
| | Accuracy | 0.625 | 0.975 | 0.700 | 0.800 | 0.875 | 0.795 |
| | Precision | 0.714 | 0.813 | 0.718 | 0.762 | 0.972 | |

Table 2: Confusion matrices, accuracies, and precisions for the network-level classifier using the uncorrected anatomically-driven deterministic connectivity.

| | | LF | LH | RF | RH | T | Total |
|---|---|---|---|---|---|---|---|
| Fully Connected Network | LF | 16 | 7 | 7 | 3 | 7 | 40 |
| | LH | 1 | 27 | 2 | 3 | 7 | 40 |
| | RF | 4 | 4 | 19 | 7 | 6 | 40 |
| | RH | 4 | 2 | 12 | 19 | 3 | 40 |
| | T | 4 | 6 | 2 | 2 | 26 | 40 |
| | Total | 29 | 46 | 42 | 34 | 49 | 200 |
| | Accuracy | 0.400 | 0.675 | 0.475 | 0.475 | 0.650 | 0.535 |
| | Precision | 0.552 | 0.587 | 0.452 | 0.559 | 0.531 | |
| Pruned Network | LF | 21 | 6 | 1 | 6 | 6 | 40 |
| | LH | 5 | 20 | 2 | 7 | 6 | 40 |
| | RF | 17 | 5 | 5 | 7 | 6 | 40 |
| | RH | 14 | 2 | 5 | 14 | 5 | 40 |
| | T | 9 | 6 | 0 | 1 | 24 | 40 |
| | Total | 66 | 39 | 13 | 35 | 47 | 200 |
| | Accuracy | 0.525 | 0.500 | 0.125 | 0.350 | 0.600 | 0.420 |
| | Precision | 0.318 | 0.513 | 0.385 | 0.400 | 0.511 | |
| Retrained Network | LF | 15 | 5 | 8 | 5 | 7 | 40 |
| | LH | 1 | 30 | 1 | 3 | 5 | 40 |
| | RF | 6 | 5 | 9 | 18 | 2 | 40 |
| | RH | 3 | 0 | 13 | 21 | 3 | 40 |
| | T | 3 | 2 | 0 | 3 | 32 | 40 |
| | Total | 28 | 42 | 31 | 50 | 49 | 200 |
| | Accuracy | 0.375 | 0.750 | 0.225 | 0.525 | 0.800 | 0.535 |
| | Precision | 0.536 | 0.714 | 0.290 | 0.420 | 0.653 | |

Table 3: Confusion matrices, accuracies, and precisions for the network-level classifier using the uncorrected population-driven deterministic connectivity with threshold 5 %.

| | | LF | LH | RF | RH | T | Total |
|---|---|---|---|---|---|---|---|
| Fully Connected Network | LF | 21 | 7 | 7 | 3 | 2 | 40 |
| | LH | 0 | 34 | 2 | 1 | 3 | 40 |
| | RF | 7 | 1 | 20 | 8 | 4 | 40 |
| | RH | 4 | 1 | 5 | 29 | 1 | 40 |
| | T | 2 | 2 | 1 | 3 | 32 | 40 |
| | Total | 34 | 45 | 35 | 44 | 42 | 200 |
| | Accuracy | 0.525 | 0.850 | 0.500 | 0.725 | 0.800 | 0.680 |
| | Precision | 0.618 | 0.756 | 0.571 | 0.659 | 0.762 | |
| Pruned Network | LF | 17 | 4 | 8 | 7 | 4 | 40 |
| | LH | 2 | 14 | 9 | 4 | 11 | 40 |
| | RF | 11 | 1 | 6 | 15 | 7 | 40 |
| | RH | 4 | 2 | 5 | 23 | 6 | 40 |
| | T | 5 | 0 | 0 | 11 | 24 | 40 |
| | Total | 39 | 21 | 28 | 60 | 52 | 200 |
| | Accuracy | 0.425 | 0.350 | 0.150 | 0.575 | 0.600 | 0.420 |
| | Precision | 0.436 | 0.667 | 0.214 | 0.383 | 0.462 | |
| Retrained Network | LF | 19 | 7 | 10 | 2 | 2 | 40 |
| | LH | 0 | 34 | 2 | 2 | 2 | 40 |
| | RF | 8 | 4 | 20 | 7 | 1 | 40 |
| | RH | 2 | 3 | 4 | 28 | 3 | 40 |
| | T | 1 | 3 | 1 | 4 | 31 | 40 |
| | Total | 30 | 51 | 37 | 43 | 39 | 200 |
| | Accuracy | 0.475 | 0.850 | 0.500 | 0.700 | 0.775 | 0.660 |
| | Precision | 0.633 | 0.667 | 0.541 | 0.651 | 0.795 | |

Table 4: Confusion matrices, accuracies, and precisions for the network-level classifier using the uncorrected population-driven deterministic connectivity with threshold 70 %.

| | | LF | LH | RF | RH | T | Total |
|---|---|---|---|---|---|---|---|
| Fully Connected Network | LF | 25 | 1 | 8 | 4 | 2 | 40 |
| | LH | 2 | 38 | 0 | 0 | 0 | 40 |
| | RF | 4 | 1 | 33 | 2 | 0 | 40 |
| | RH | 1 | 1 | 2 | 34 | 2 | 40 |
| | T | 1 | 1 | 0 | 3 | 35 | 40 |
| | Total | 33 | 42 | 43 | 43 | 39 | 200 |
| | Accuracy | 0.625 | 0.950 | 0.825 | 0.850 | 0.875 | 0.825 |
| | Precision | 0.758 | 0.905 | 0.767 | 0.791 | 0.897 | |
| Pruned Network | LF | 14 | 11 | 9 | 2 | 4 | 40 |
| | LH | 1 | 24 | 13 | 1 | 1 | 40 |
| | RF | 3 | 3 | 18 | 1 | 15 | 40 |
| | RH | 3 | 10 | 8 | 12 | 7 | 40 |
| | T | 2 | 8 | 9 | 6 | 15 | 40 |
| | Total | 23 | 56 | 57 | 22 | 42 | 200 |
| | Accuracy | 0.350 | 0.600 | 0.450 | 0.300 | 0.375 | 0.415 |
| | Precision | 0.609 | 0.429 | 0.316 | 0.545 | 0.357 | |
| Retrained Network | LF | 22 | 3 | 9 | 2 | 4 | 40 |
| | LH | 1 | 39 | 0 | 0 | 0 | 40 |
| | RF | 0 | 0 | 34 | 3 | 3 | 40 |
| | RH | 1 | 3 | 6 | 29 | 1 | 40 |
| | T | 2 | 0 | 2 | 3 | 33 | 40 |
| | Total | 26 | 45 | 51 | 37 | 41 | 200 |
| | Accuracy | 0.550 | 0.975 | 0.850 | 0.725 | 0.825 | 0.785 |
| | Precision | 0.846 | 0.867 | 0.667 | 0.784 | 0.805 | |

Table 5: Confusion matrices, accuracies, and precisions for the network-level classifier using the uncorrected population-driven deterministic connectivity with threshold 99 %.

| | | LF | LH | RF | RH | T | Total |
|---|---|---|---|---|---|---|---|
| Fully Connected Network | LF | 5 | 14 | 21 | 0 | 0 | 40 |
| | LH | 3 | 31 | 5 | 1 | 0 | 40 |
| | RF | 1 | 14 | 24 | 0 | 1 | 40 |
| | RH | 0 | 13 | 25 | 2 | 0 | 40 |
| | T | 1 | 21 | 17 | 0 | 1 | 40 |
| | Total | 10 | 93 | 92 | 3 | 2 | 200 |
| | Accuracy | 0.125 | 0.775 | 0.600 | 0.050 | 0.025 | 0.315 |
| | Precision | 0.500 | 0.333 | 0.261 | 0.667 | 0.500 | |
| Pruned Network | LF | 24 | 13 | 1 | 0 | 2 | 40 |
| | LH | 9 | 30 | 0 | 0 | 1 | 40 |
| | RF | 25 | 14 | 0 | 0 | 1 | 40 |
| | RH | 26 | 13 | 0 | 1 | 0 | 40 |
| | T | 17 | 21 | 1 | 0 | 1 | 40 |
| | Total | 101 | 91 | 2 | 1 | 5 | 200 |
| | Accuracy | 0.600 | 0.750 | 0.000 | 0.025 | 0.025 | 0.280 |
| | Precision | 0.238 | 0.330 | 0.000 | 1.000 | 0.200 | |
| Retrained Network | LF | 3 | 13 | 22 | 0 | 2 | 40 |
| | LH | 3 | 31 | 6 | 0 | 0 | 40 |
| | RF | 0 | 14 | 25 | 0 | 1 | 40 |
| | RH | 0 | 13 | 26 | 1 | 0 | 40 |
| | T | 2 | 21 | 16 | 0 | 1 | 40 |
| | Total | 8 | 92 | 95 | 1 | 4 | 200 |
| | Accuracy | 0.075 | 0.775 | 0.625 | 0.025 | 0.025 | 0.305 |
| | Precision | 0.375 | 0.337 | 0.263 | 1.000 | 0.250 | |

Table 6: Confusion matrices, accuracies, and precisions for the network-level classifier using the uncorrected probabilistic connectivity with threshold −4.

| | | LF | LH | RF | RH | T | Total |
|---|---|---|---|---|---|---|---|
| Fully Connected Network | LF | 5 | 7 | 22 | 3 | 3 | 40 |
| | LH | 1 | 27 | 5 | 6 | 1 | 40 |
| | RF | 2 | 8 | 24 | 4 | 2 | 40 |
| | RH | 3 | 7 | 10 | 15 | 5 | 40 |
| | T | 2 | 19 | 4 | 8 | 7 | 40 |
| | Total | 13 | 68 | 65 | 36 | 18 | 200 |
| | Accuracy | 0.125 | 0.675 | 0.600 | 0.375 | 0.175 | 0.390 |
| | Precision | 0.385 | 0.397 | 0.369 | 0.417 | 0.389 | |
| Pruned Network | LF | 11 | 0 | 16 | 7 | 6 | 40 |
| | LH | 5 | 5 | 3 | 5 | 22 | 40 |
| | RF | 8 | 0 | 19 | 6 | 7 | 40 |
| | RH | 5 | 4 | 7 | 18 | 6 | 40 |
| | T | 3 | 2 | 2 | 13 | 20 | 40 |
| | Total | 32 | 11 | 47 | 49 | 61 | 200 |
| | Accuracy | 0.275 | 0.125 | 0.475 | 0.450 | 0.500 | 0.365 |
| | Precision | 0.344 | 0.455 | 0.404 | 0.367 | 0.328 | |
| Retrained Network | LF | 6 | 5 | 22 | 5 | 2 | 40 |
| | LH | 0 | 28 | 3 | 7 | 2 | 40 |
| | RF | 2 | 5 | 26 | 5 | 2 | 40 |
| | RH | 1 | 8 | 10 | 20 | 1 | 40 |
| | T | 1 | 16 | 6 | 9 | 8 | 40 |
| | Total | 10 | 62 | 67 | 46 | 15 | 200 |
| | Accuracy | 0.150 | 0.700 | 0.650 | 0.500 | 0.200 | 0.440 |
| | Precision | 0.600 | 0.452 | 0.388 | 0.435 | 0.533 | |

Table 7: Confusion matrices, accuracies, and precisions for the network-level classifier using the uncorrected probabilistic connectivity with threshold $-3$.

| | | LF | LH | RF | RH | T | Total |
|---|---|---|---|---|---|---|---|
| Fully Connected Network | LF | 24 | 1 | 10 | 4 | 1 | 40 |
| | LH | 0 | 39 | 1 | 0 | 0 | 40 |
| | RF | 7 | 2 | 26 | 4 | 1 | 40 |
| | RH | 0 | 2 | 9 | 29 | 0 | 40 |
| | T | 1 | 0 | 2 | 2 | 35 | 40 |
| | Total | 32 | 44 | 48 | 39 | 37 | 200 |
| | Accuracy | 0.600 | 0.975 | 0.650 | 0.725 | 0.875 | 0.765 |
| | Precision | 0.750 | 0.886 | 0.542 | 0.744 | 0.946 | |
| Pruned Network | LF | 21 | 3 | 8 | 7 | 1 | 40 |
| | LH | 13 | 19 | 1 | 3 | 4 | 40 |
| | RF | 10 | 2 | 18 | 10 | 0 | 40 |
| | RH | 4 | 1 | 1 | 34 | 0 | 40 |
| | T | 2 | 2 | 0 | 12 | 24 | 40 |
| | Total | 50 | 27 | 28 | 66 | 29 | 200 |
| | Accuracy | 0.525 | 0.475 | 0.450 | 0.850 | 0.600 | 0.580 |
| | Precision | 0.420 | 0.704 | 0.643 | 0.515 | 0.828 | |
| Retrained Network | LF | 23 | 1 | 13 | 2 | 1 | 40 |
| | LH | 1 | 35 | 1 | 2 | 1 | 40 |
| | RF | 5 | 2 | 28 | 4 | 1 | 40 |
| | RH | 0 | 2 | 8 | 29 | 1 | 40 |
| | T | 1 | 1 | 3 | 0 | 35 | 40 |
| | Total | 30 | 41 | 53 | 37 | 39 | 200 |
| | Accuracy | 0.575 | 0.875 | 0.700 | 0.725 | 0.875 | 0.750 |
| | Precision | 0.767 | 0.854 | 0.528 | 0.784 | 0.897 | |

Table 8: Confusion matrices, accuracies, and precisions for the network-level classifier using the uncorrected probabilistic connectivity with threshold −2.

| | | LF | LH | RF | RH | T | Total |
|---|---|---|---|---|---|---|---|
| Fully Connected Network | LF | 27 | 0 | 7 | 3 | 3 | 40 |
| | LH | 1 | 38 | 1 | 0 | 0 | 40 |
| | RF | 4 | 1 | 31 | 3 | 1 | 40 |
| | RH | 0 | 3 | 2 | 35 | 0 | 40 |
| | T | 2 | 0 | 0 | 3 | 35 | 40 |
| | Total | 34 | 42 | 41 | 44 | 39 | 200 |
| | Accuracy | 0.675 | 0.950 | 0.775 | 0.875 | 0.875 | 0.830 |
| | Precision | 0.794 | 0.905 | 0.756 | 0.795 | 0.897 | |
| Pruned Network | LF | 29 | 1 | 6 | 3 | 1 | 40 |
| | LH | 5 | 34 | 1 | 0 | 0 | 40 |
| | RF | 7 | 2 | 28 | 3 | 0 | 40 |
| | RH | 1 | 5 | 3 | 31 | 0 | 40 |
| | T | 2 | 0 | 1 | 2 | 35 | 40 |
| | Total | 44 | 42 | 39 | 39 | 36 | 200 |
| | Accuracy | 0.725 | 0.850 | 0.700 | 0.775 | 0.875 | 0.785 |
| | Precision | 0.659 | 0.810 | 0.718 | 0.795 | 0.972 | |
| Retrained Network | LF | 27 | 1 | 6 | 4 | 2 | 40 |
| | LH | 0 | 39 | 0 | 1 | 0 | 40 |
| | RF | 7 | 1 | 29 | 2 | 1 | 40 |
| | RH | 1 | 4 | 2 | 33 | 0 | 40 |
| | T | 2 | 0 | 0 | 1 | 37 | 40 |
| | Total | 37 | 45 | 37 | 41 | 40 | 200 |
| | Accuracy | 0.675 | 0.975 | 0.725 | 0.825 | 0.925 | 0.825 |
| | Precision | 0.730 | 0.867 | 0.784 | 0.805 | 0.925 | |

Table 9: Confusion matrices, accuracies, and precisions for the network-level classifier using the corrected anatomically-driven deterministic connectivity.

| | | LF | LH | RF | RH | T | Total |
|---|---|---|---|---|---|---|---|
| Fully Connected Network | LF | 21 | 6 | 5 | 3 | 5 | 40 |
| | LH | 3 | 36 | 1 | 0 | 0 | 40 |
| | RF | 3 | 4 | 26 | 4 | 3 | 40 |
| | RH | 3 | 1 | 9 | 23 | 4 | 40 |
| | T | 4 | 1 | 1 | 4 | 30 | 40 |
| | Total | 34 | 48 | 42 | 34 | 42 | 200 |
| | Accuracy | 0.525 | 0.900 | 0.650 | 0.575 | 0.750 | 0.680 |
| | Precision | 0.618 | 0.750 | 0.619 | 0.676 | 0.714 | |
| Pruned Network | LF | 13 | 5 | 11 | 6 | 5 | 40 |
| | LH | 16 | 10 | 3 | 0 | 11 | 40 |
| | RF | 8 | 2 | 10 | 13 | 7 | 40 |
| | RH | 2 | 3 | 9 | 17 | 9 | 40 |
| | T | 1 | 3 | 3 | 10 | 23 | 40 |
| | Total | 40 | 23 | 36 | 46 | 55 | 200 |
| | Accuracy | 0.325 | 0.250 | 0.250 | 0.425 | 0.575 | 0.365 |
| | Precision | 0.325 | 0.435 | 0.278 | 0.370 | 0.418 | |
| Retrained Network | LF | 20 | 6 | 7 | 3 | 4 | 40 |
| | LH | 2 | 35 | 2 | 1 | 0 | 40 |
| | RF | 3 | 4 | 27 | 3 | 3 | 40 |
| | RH | 0 | 4 | 12 | 21 | 3 | 40 |
| | T | 3 | 2 | 1 | 2 | 32 | 40 |
| | Total | 28 | 51 | 49 | 30 | 42 | 200 |
| | Accuracy | 0.500 | 0.875 | 0.675 | 0.525 | 0.800 | 0.675 |
| | Precision | 0.714 | 0.686 | 0.551 | 0.700 | 0.762 | |

Table 10: Confusion matrices, accuracies, and precisions for the network-level classifier using the corrected population-driven deterministic connectivity with threshold 5 %.

| | | LF | LH | RF | RH | T | Total |
|---|---|---|---|---|---|---|---|
| Fully Connected Network | LF | 22 | 2 | 10 | 2 | 4 | 40 |
| | LH | 2 | 28 | 3 | 1 | 6 | 40 |
| | RF | 7 | 3 | 19 | 7 | 4 | 40 |
| | RH | 0 | 3 | 4 | 31 | 2 | 40 |
| | T | 2 | 1 | 2 | 5 | 30 | 40 |
| | Total | 33 | 37 | 38 | 46 | 46 | 200 |
| | Accuracy | 0.550 | 0.700 | 0.475 | 0.775 | 0.750 | 0.650 |
| | Precision | 0.667 | 0.757 | 0.500 | 0.674 | 0.652 | |
| Pruned Network | LF | 21 | 9 | 5 | 3 | 2 | 40 |
| | LH | 4 | 15 | 6 | 10 | 5 | 40 |
| | RF | 9 | 6 | 12 | 4 | 9 | 40 |
| | RH | 6 | 4 | 4 | 19 | 7 | 40 |
| | T | 2 | 2 | 1 | 15 | 20 | 40 |
| | Total | 42 | 36 | 28 | 51 | 43 | 200 |
| | Accuracy | 0.525 | 0.375 | 0.300 | 0.475 | 0.500 | 0.435 |
| | Precision | 0.500 | 0.417 | 0.429 | 0.373 | 0.465 | |
| Retrained Network | LF | 17 | 4 | 9 | 4 | 6 | 40 |
| | LH | 3 | 31 | 3 | 1 | 2 | 40 |
| | RF | 7 | 4 | 15 | 9 | 5 | 40 |
| | RH | 1 | 4 | 6 | 26 | 3 | 40 |
| | T | 1 | 1 | 5 | 2 | 31 | 40 |
| | Total | 29 | 44 | 38 | 42 | 47 | 200 |
| | Accuracy | 0.425 | 0.775 | 0.375 | 0.650 | 0.775 | 0.600 |
| | Precision | 0.586 | 0.705 | 0.395 | 0.619 | 0.660 | |

Table 11: Confusion matrices, accuracies, and precisions for the network-level classifier using the corrected population-driven deterministic connectivity with threshold 70 %.

| | | LF | LH | RF | RH | T | Total |
|---|---|---|---|---|---|---|---|
| Fully Connected Network | LF | 24 | 4 | 9 | 0 | 3 | 40 |
| | LH | 1 | 37 | 0 | 1 | 1 | 40 |
| | RF | 3 | 0 | 30 | 4 | 3 | 40 |
| | RH | 1 | 4 | 4 | 30 | 1 | 40 |
| | T | 1 | 0 | 0 | 3 | 36 | 40 |
| | Total | 30 | 45 | 43 | 38 | 44 | 200 |
| | Accuracy | 0.600 | 0.925 | 0.750 | 0.750 | 0.900 | 0.785 |
| | Precision | 0.800 | 0.822 | 0.698 | 0.789 | 0.818 | |
| Pruned Network | LF | 30 | 0 | 5 | 0 | 5 | 40 |
| | LH | 1 | 36 | 0 | 2 | 1 | 40 |
| | RF | 8 | 0 | 22 | 6 | 4 | 40 |
| | RH | 5 | 6 | 4 | 24 | 1 | 40 |
| | T | 3 | 1 | 1 | 2 | 33 | 40 |
| | Total | 47 | 43 | 32 | 34 | 44 | 200 |
| | Accuracy | 0.750 | 0.900 | 0.550 | 0.600 | 0.825 | 0.725 |
| | Precision | 0.638 | 0.837 | 0.688 | 0.706 | 0.750 | |
| Retrained Network | LF | 25 | 2 | 8 | 1 | 4 | 40 |
| | LH | 1 | 35 | 0 | 2 | 2 | 40 |
| | RF | 4 | 0 | 29 | 5 | 2 | 40 |
| | RH | 1 | 6 | 2 | 29 | 2 | 40 |
| | T | 2 | 0 | 0 | 1 | 37 | 40 |
| | Total | 33 | 43 | 39 | 38 | 47 | 200 |
| | Accuracy | 0.625 | 0.875 | 0.725 | 0.725 | 0.925 | 0.775 |
| | Precision | 0.758 | 0.814 | 0.744 | 0.763 | 0.787 | |

Table 12: Confusion matrices, accuracies, and precisions for the network-level classifier using the corrected population-driven deterministic connectivity with threshold 99 %.

| | | LF | LH | RF | RH | T | Total |
|---|---|---|---|---|---|---|---|
| Fully Connected Network | LF | 25 | 2 | 9 | 1 | 3 | 40 |
| | LH | 4 | 31 | 0 | 2 | 3 | 40 |
| | RF | 6 | 2 | 27 | 4 | 1 | 40 |
| | RH | 4 | 2 | 4 | 30 | 0 | 40 |
| | T | 3 | 2 | 4 | 3 | 28 | 40 |
| | Total | 42 | 39 | 44 | 40 | 35 | 200 |
| | Accuracy | 0.625 | 0.775 | 0.675 | 0.750 | 0.700 | 0.705 |
| | Precision | 0.595 | 0.795 | 0.614 | 0.750 | 0.800 | |
| Pruned Network | LF | 17 | 1 | 5 | 16 | 1 | 40 |
| | LH | 22 | 11 | 0 | 6 | 1 | 40 |
| | RF | 14 | 1 | 5 | 20 | 0 | 40 |
| | RH | 7 | 3 | 1 | 27 | 2 | 40 |
| | T | 18 | 1 | 2 | 14 | 5 | 40 |
| | Total | 78 | 17 | 13 | 83 | 9 | 200 |
| | Accuracy | 0.425 | 0.275 | 0.125 | 0.675 | 0.125 | 0.325 |
| | Precision | 0.218 | 0.647 | 0.385 | 0.325 | 0.556 | |
| Retrained Network | LF | 26 | 3 | 8 | 2 | 1 | 40 |
| | LH | 5 | 32 | 0 | 1 | 2 | 40 |
| | RF | 8 | 3 | 24 | 5 | 0 | 40 |
| | RH | 2 | 2 | 5 | 29 | 2 | 40 |
| | T | 3 | 1 | 3 | 5 | 28 | 40 |
| | Total | 44 | 41 | 40 | 42 | 33 | 200 |
| | Accuracy | 0.650 | 0.800 | 0.600 | 0.725 | 0.700 | 0.695 |
| | Precision | 0.591 | 0.780 | 0.600 | 0.690 | 0.848 | |

Table 13: Confusion matrices, accuracies, and precisions for the network-level classifier using the corrected probabilistic connectivity with threshold −4.

| | | LF | LH | RF | RH | T | Total |
|---|---|---|---|---|---|---|---|
| Fully Connected Network | LF | 21 | 2 | 15 | 2 | 0 | 40 |
| | LH | 1 | 37 | 1 | 0 | 1 | 40 |
| | RF | 8 | 2 | 26 | 4 | 0 | 40 |
| | RH | 0 | 1 | 3 | 36 | 0 | 40 |
| | T | 2 | 0 | 1 | 3 | 34 | 40 |
| | Total | 32 | 42 | 46 | 45 | 35 | 200 |
| | Accuracy | 0.525 | 0.925 | 0.650 | 0.900 | 0.850 | 0.770 |
| | Precision | 0.656 | 0.881 | 0.565 | 0.800 | 0.971 | |
| Pruned Network | LF | 16 | 10 | 2 | 9 | 3 | 40 |
| | LH | 3 | 33 | 0 | 3 | 1 | 40 |
| | RF | 5 | 10 | 7 | 14 | 4 | 40 |
| | RH | 0 | 7 | 2 | 29 | 2 | 40 |
| | T | 3 | 18 | 0 | 10 | 9 | 40 |
| | Total | 27 | 78 | 11 | 65 | 19 | 200 |
| | Accuracy | 0.400 | 0.825 | 0.175 | 0.725 | 0.225 | 0.470 |
| | Precision | 0.593 | 0.423 | 0.636 | 0.446 | 0.474 | |
| Retrained Network | LF | 22 | 4 | 12 | 2 | 0 | 40 |
| | LH | 0 | 39 | 0 | 0 | 1 | 40 |
| | RF | 7 | 2 | 27 | 4 | 0 | 40 |
| | RH | 0 | 1 | 5 | 32 | 2 | 40 |
| | T | 2 | 0 | 2 | 5 | 31 | 40 |
| | Total | 31 | 46 | 46 | 43 | 34 | 200 |
| | Accuracy | 0.550 | 0.975 | 0.675 | 0.800 | 0.775 | 0.755 |
| | Precision | 0.710 | 0.848 | 0.587 | 0.744 | 0.912 | |

Table 14: Confusion matrices, accuracies, and precisions for the network-level classifier using the corrected probabilistic connectivity with threshold −3.

| | | LF | LH | RF | RH | T | Total |
|---|---|---|---|---|---|---|---|
| Fully Connected Network | LF | 27 | 1 | 9 | 2 | 1 | 40 |
| | LH | 1 | 37 | 1 | 1 | 0 | 40 |
| | RF | 4 | 1 | 31 | 3 | 1 | 40 |
| | RH | 0 | 1 | 4 | 35 | 0 | 40 |
| | T | 1 | 1 | 2 | 4 | 32 | 40 |
| | Total | 33 | 41 | 47 | 45 | 34 | 200 |
| | Accuracy | 0.675 | 0.925 | 0.775 | 0.875 | 0.800 | 0.810 |
| | Precision | 0.818 | 0.902 | 0.660 | 0.778 | 0.941 | |
| Pruned Network | LF | 19 | 1 | 12 | 7 | 1 | 40 |
| | LH | 0 | 34 | 2 | 2 | 2 | 40 |
| | RF | 2 | 3 | 25 | 9 | 1 | 40 |
| | RH | 1 | 2 | 11 | 26 | 0 | 40 |
| | T | 3 | 4 | 2 | 12 | 19 | 40 |
| | Total | 25 | 44 | 52 | 56 | 23 | 200 |
| | Accuracy | 0.475 | 0.850 | 0.625 | 0.650 | 0.475 | 0.615 |
| | Precision | 0.760 | 0.773 | 0.481 | 0.464 | 0.826 | |
| Retrained Network | LF | 25 | 2 | 10 | 2 | 1 | 40 |
| | LH | 2 | 37 | 0 | 1 | 0 | 40 |
| | RF | 3 | 1 | 32 | 3 | 1 | 40 |
| | RH | 0 | 1 | 4 | 35 | 0 | 40 |
| | T | 2 | 0 | 1 | 4 | 33 | 40 |
| | Total | 32 | 41 | 47 | 45 | 35 | 200 |
| | Accuracy | 0.625 | 0.925 | 0.800 | 0.875 | 0.825 | 0.810 |
| | Precision | 0.781 | 0.902 | 0.681 | 0.778 | 0.943 | |

Table 15: Confusion matrices, accuracies, and precisions for the network-level classifier using the corrected probabilistic connectivity with threshold −2.
